# Development of a NanoBiT based high throughput screening assay for discovery of NOS1-CAPON interaction inhibitors

**DOI:** 10.64898/2026.01.25.701608

**Authors:** Sungwoo Cho, Moustafa T. Gabr

**Affiliations:** Department of Radiology, Molecular Imaging Innovations Institute (MI3), Weill Cornell Medicine, New York, NY 10065, USA

**Keywords:** NOS1, CAPON, NanoBiT, high-throughput screening, protein-protein interaction, drug discovery

## Abstract

The interaction between neuronal nitric oxide synthase (NOS1) and its adaptor protein CAPON (NOS1AP) plays a critical role in various neurological processes and has been implicated in cardiovascular and neuropsychiatric disorders. Disruption of this protein-protein interaction represents a potential therapeutic strategy, yet identifying small molecule inhibitors has been challenging. Here, we present the development and validation of a NanoBiT-based luminescence complementation assay optimized for high-throughput screening (HTS) of NOS1-NOS1AP interaction inhibitors. We engineered NOS1 and NOS1AP fusion proteins with HiBiT and LgBiT complementary subunits, respectively, and established stable CHO-K1 cell lines for robust signal generation. The assay demonstrated excellent performance characteristics with a signal-to-background ratio exceeding 240-fold, and was validated using TAT-GESV, a known peptide inhibitor that showed time- and dose-dependent inhibition. We successfully screened a diverse library of 10,240 compounds and identified 19 validated hits with IC50 values ranging from 2.54 to greater than 30 μM, with the majority exhibiting IC_50_ values below 30 μM. The top three compounds exhibited potent inhibitory activity with IC_50_ values of less than 5 μM. This NanoBiT-based assay provides a reliable and efficient platform for discovering novel NOS1-NOS1AP interaction inhibitors and can be adapted for other protein-protein interaction studies.

## Introduction

Neuronal nitric oxide synthase (NOS1, also known as nNOS) is a critical enzyme responsible for producing nitric oxide (NO) in neurons, playing essential roles in neurotransmission, synaptic plasticity, and neuronal development [1, 2]. NOS1 activity is tightly regulated through protein-protein interactions, with NOS1-associated protein (NOS1AP, also known as CAPON) serving as a key adaptor that modulates NOS1 localization and function [3]. NOS1AP was first identified by Jaffrey et al. as a protein that interacts with the PDZ domain of NOS1 through its C-terminal PDZ-binding motif [3]. This interaction competes with PSD-95 for NOS1 binding, thereby regulating NOS1’s association with NMDA receptor complexes and affecting downstream signaling pathways [4].

The NOS1-NOS1AP interaction has been implicated in diverse pathological conditions. Genetic variants in NOS1AP have been associated with QT interval prolongation and increased risk of sudden cardiac death in multiple population studies [5-7]. In the central nervous system, NOS1AP has emerged as a susceptibility gene for schizophrenia and other psychiatric disorders, with increased expression observed in postmortem brain samples from affected individuals [8-10]. Additionally, the NOS1-NOS1AP pathway has been linked to excitotoxic neuronal death, making it an attractive therapeutic target for neuroprotection [11]. These diverse disease associations underscore the therapeutic potential of modulating this protein-protein interaction.

Despite the therapeutic potential, identifying small molecule inhibitors of the NOS1-NOS1AP interaction has been challenging due to the large binding interface and the lack of suitable high-throughput screening assays. Traditional methods for detecting protein-protein interactions, such as co-immunoprecipitation and yeast two-hybrid systems, are labor-intensive and not amenable to large-scale compound screening [12]. Fluorescence-based assays like FRET and BRET offer improved throughput but often suffer from limited dynamic range and complex signal processing requirements [13].

The NanoBiT (NanoLuc Binary Technology) system represents a significant advancement in protein complementation assays, offering superior brightness, stability, and signal-to-noise ratio compared to traditional methods [14]. NanoBiT is based on the structural complementation of two subunits derived from NanoLuc luciferase: a large subunit (LgBiT, 18 kDa) and small subunits of varying affinities. The original system employed SmBiT (11 amino acids) with low affinity (K_d_ = 190 μM) to LgBiT, ensuring that complementation is driven by the interaction of the target proteins rather than intrinsic subunit affinity [14]. Subsequently, Schwinn et al. identified HiBiT, a high-affinity variant (K_d_ = 700 pM) that enables efficient complementation and serves as a quantitative luminescent tag for protein detection [15]. The HiBiT system offers several advantages including bright luminescence over an 8-order magnitude linear range and minimal interference with protein function due to its small size [15, 16]. When proteins of interest fused to these subunits interact, the NanoLuc enzyme is reconstituted, producing a bright and stable luminescent signal [14]. This technology has been successfully applied to study various protein-protein interactions and has proven particularly valuable for drug discovery applications [17, 18].

In this study, we developed and validated a NanoBiT-based assay specifically designed for high-throughput screening of NOS1-NOS1AP interaction inhibitors. We optimized the assay conditions, validated its performance using a known peptide inhibitor (TAT-GESV), and successfully screened a diverse chemical library to identify novel small molecule hits. Our results demonstrate that this assay provides a robust and efficient platform for discovering NOS1-NOS1AP interaction inhibitors with potential therapeutic applications.

## Methods

### Construction of NanoBiT tagged proteins

Based on the well-characterized interaction interface between NOS1 and NOS1AP, where the C-terminal PDZ-binding motif of NOS1AP binds to the N-terminal PDZ domain of NOS1 [3, 4], we designed our NanoBiT constructs to preserve this interaction while enabling efficient complementation. NOS1AP was fused to LgBiT at its C-terminus through a flexible linker peptide (SSGGGGGS), creating the NOS1AP-LgBiT construct. This orientation positions the LgBiT subunit adjacent to the PDZ-binding domain of NOS1AP while allowing flexibility for interaction. NOS1 was fused to HiBiT at its N-terminus using a long flexible linker (GNSGSSGGGGSGGGGSSG) to minimize steric interference with the PDZ domain, generating the HiBiT-NOS1 construct. HiBiT was selected over SmBiT for its high affinity (K_d_ = 700 pM) to LgBiT, which ensures robust signal generation even at low expression levels typical of stable cell lines [15]. Both constructs were cloned into pcDNA3.1 mammalian expression vectors under the control of a CMV promoter using standard molecular cloning techniques.

### Cell culture and stable cell line generation

CHO-K1 cells (Chinese Hamster Ovary; ATCC CCL-61) were cultured in RPMI-1640 medium (Thermofisher Scienctific, MA, USA) supplemented with 10% fetal bovine serum and 1% penicillin-streptomycin (100 U/mL penicillin, 100 μg/mL streptomycin) at 37°C in a humidified atmosphere containing 5% CO_2_. Stable cell lines co-expressing HiBiT-NOS1 and NOS1AP-LgBiT were generated by co-transfection using Lipofectamine 3000 (Thermofisher Scienctific, MA, USA) according to the manufacturer’s instructions, followed by selection with 10 μg/mL puromycin and 20 μg/mL blastacidin for 2-3 weeks. Single clones were isolated and screened for optimal NanoBiT signal intensity and signal-to-background ratio. The selected clone demonstrating the highest signal-to-background ratio was expanded.

### NanoBiT assay protocol

CHO-K1 cells stably expressing HiBiT-NOS1 and NOS1AP-LgBiT were seeded in white 96-well plates at a density of 2 × 10^4^ cells per well and cultured overnight to achieve approximately 80% confluency. For compound treatment, the culture medium was carefully aspirated and replaced with 100 μL of Hank’s Balanced Salt Solution (HBSS). Test compounds (dissolved in DMSO), TAT-GESV peptide, or vehicle controls were added at the indicated concentrations in 100 μL volume and incubated at 37°C for indicated hours. After incubation, 100 μL of HBSS containing 20 μM furimazine (AOBIOUS, Gloucester, MA, USA) was added to each well, achieving a final furimazine concentration of 10 μM. The plates were then incubated at 37°C for 10 minutes to allow substrate equilibration. Following a 10minute, luminescence was measured using a Tecan Spark multimode microplate reader with a spectral range of 450-480 nm (emission peak at 460 nm).

### TAT-GESV Peptide validation

TAT-GESV is a cell-penetrating peptide designed to disrupt NOS1-NOS1AP interaction by competing for the PDZ domain binding site of NOS1 [11]. The peptide contains the HIV TAT cell-penetrating sequence fused to the C-terminal GESV motif derived from NOS1AP, which specifically binds to the unique PDZ domain of NOS1 [11, 19]. TAT-GESV peptide (>95% purity; synthesized by in-house) was dissolved in sterile water to prepare a 1 mM stock solution. To validate the NanoBiT assay system, cells were treated with various concentrations of TAT-GESV (0.1, 0.3, 1, 3, 10, and 30 μM) for different time periods (15 minutes, 1 hour, and 3 hours) to assess dose-response and time-dependent effects. The NanoBiT signal was measured as described above and expressed as a percentage of vehicle control. PBS was used as a negative control.

### Hight throughput screening campaign

For the HTS campaign, compounds from the Discovery Diversity Set (DDS) library (Enamine, NJ, USA) were reformatted into a Discovery Diversity Set Daughter (DDSD) library and screened in a 4-in-1 pooled format, where four structurally diverse compounds were combined in each well at a final concentration of 25 μM per compound. This pooling strategy enables efficient primary screening while maintaining sufficient compound concentration for hit identification. The screening was performed in white 96-well plates. Hit compounds were defined as those showing ≥60% inhibition of the NanoBiT signal relative to the vehicle control.

### Statical analysis

All data are presented as mean ± standard deviation (SD) from at least three independent experiments. Statistical analyses were performed using GraphPad Prism 9.0 software.

## Results

### Establishment and validation of the NanoBiT based assay system

To develop a robust screening platform for NOS1-NOS1AP interaction inhibitors, we designed and constructed NanoBiT-tagged proteins based on the well-established interaction interface between these proteins. Previous structural and biochemical studies have demonstrated that the C-terminal PDZ-binding motif (EIAV/GESV) of NOS1AP specifically interacts with the N-terminal PDZ domain of NOS1, and this interaction is essential for the regulatory function of NOS1AP [3, 19]. Accordingly, we designed our constructs to preserve this interaction while enabling efficient NanoBiT complementation: NOS1AP was tagged with LgBiT at the C-terminus, while NOS1 was tagged with HiBiT at the N-terminus using appropriate flexible linker sequences (Figure 1A). The principle of the assay is depicted in Figure 1B: when NOS1 and NOS1AP interact, the HiBiT and LgBiT subunits are brought into proximity, enabling NanoLuciferase complementation and generating a bright luminescent signal at 460 nm. Conversely, when an inhibitor disrupts this interaction, the complementation is prevented, resulting in reduced luminescence.

**Fig 1.**
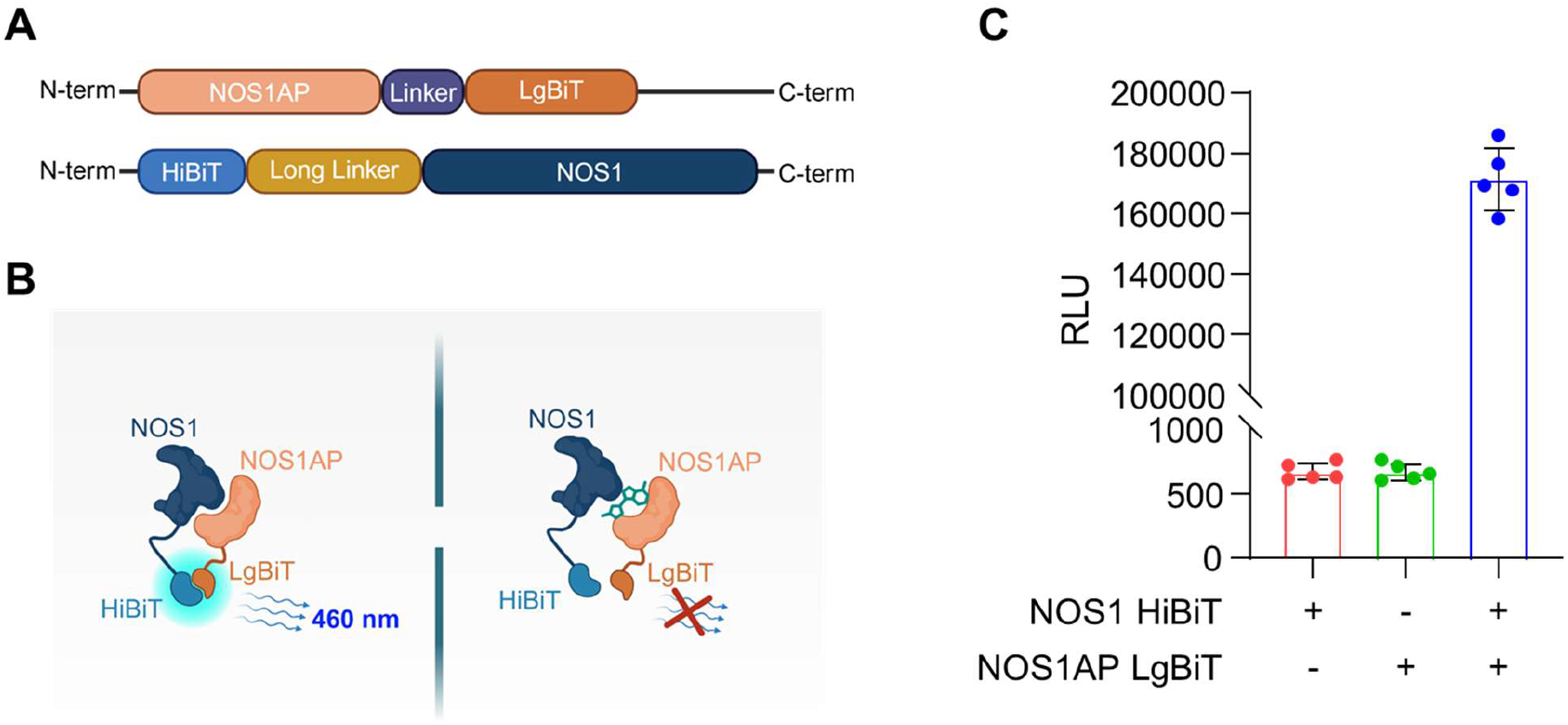
Establishment of NanoBiT-based assay system for monitoring NOS1-NOS1AP interaction. (A) Schematic representation of NanoBiT constructs. NOS1AP was tagged to LgBiT at the C-terminus, and NOS1 was tagged to HiBiT at the N-terminus. (B) Principle of NanoBiT assay. Left: NOS1-NOS1AP interaction brings HiBiT and LgBiT into proximity, enabling NanoLuciferase complementation and light emission. Right: Inhibitor disrupts the interaction, preventing luminescent signal generation. (C) Validation of the NanoBiT system in CHO-K1 cells. Luciferase luminescence measured upon expression of each tagged protein in CHO-K1 cells. Data are presented as mean ± SD (n = 5).

We established stable CHO-K1 cell lines co-expressing both HiBiT-NOS1 and NOS1AP-LgBiT constructs and validated the system by measuring luminescence upon expression of individual components versus both proteins together (Figure 1C). Expression of HiBiT-NOS1 alone or NOS1AP-LgBiT alone produced minimal background signal (approximately 600-700 RLU). In contrast, co-expression of both fusion proteins resulted in a dramatic increase in luminescence to approximately 170,000 RLU, representing a signal-to-background ratio of greater than 240-fold. This robust signal generation confirmed successful reconstitution of NanoLuciferase activity through NOS1-NOS1AP interaction and demonstrated the suitability of this system for HTS applications.

### Validation with TAT-GESV Peptide Inhibitor

To confirm that the NanoBiT assay could detect inhibition of the NOS1-NOS1AP interaction, we tested TAT-GESV, a cell-penetrating peptide previously demonstrated to compete for the nNOS PDZ domain and disrupt NOS1-NOS1AP interaction in cortical neurons [11]. The researchers showed that TAT-GESV inhibits the recruitment of NOS1AP to NOS1 following excitotoxic stimulus and provides neuroprotection in models of hypoxia-ischemia, validating this peptide as an inhibitor of the NOS1-NOS1AP interaction [11].

We evaluated the time-dependence of inhibition by treating cells with various concentrations of TAT-GESV for 15 minutes, 1 hour, or 3 hours (Figure 2A-C). At 15 minutes, TAT-GESV showed minimal inhibitory effect even at the highest concentration tested (30 μM), with NanoBiT activity remaining above 90% of control (Figure 2A). After 1 hour of treatment, modest inhibition was observed, with activity decreasing to approximately 90% at 30 μM (Figure 2B). The most pronounced inhibition occurred after 3 hours of treatment, where TAT-GESV demonstrated clear dose-dependent inhibition with activity reduced to approximately 55% at 30 μM concentration (Figure 2C). These time-dependent inhibition profiles are consistent with the known mechanism of TAT-mediated peptide delivery, which requires cellular uptake, endosomal escape, and equilibration with intracellular targets. The results confirmed that the NanoBiT assay system responds appropriately to a known NOS1-NOS1AP interaction inhibitor and established that an optimal compound incubation time of 3 hours should be used for the screening campaign.

**Figure 2.**
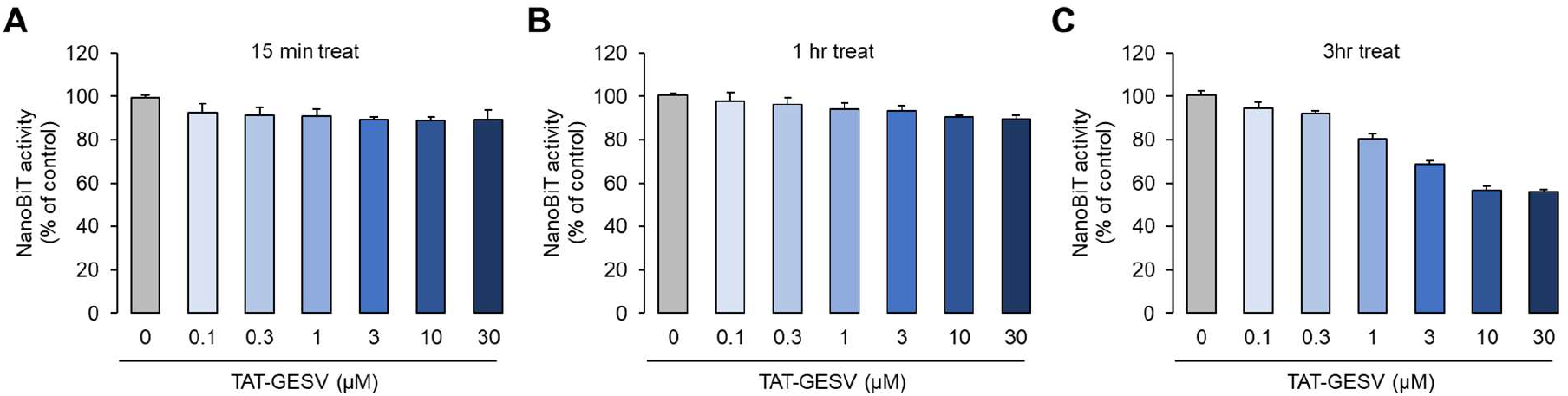
Validation of NanoBiT assay system using TAT-GESV peptide. CHO-K1 cells stably expressing NOS1-HiBiT and NOS1AP-LgBiT were treated with indicated concentrations of TAT-GESV peptide for (A) 15 minutes, (B) 1 hour, and (C) 3 hours. NanoBiT signal was measured and expressed as a percentage of the vehicle control. TAT-GESV showed a time- and dose-dependent inhibition of NOS1-NOS1AP interaction, with the most pronounced effect observed at 3 hours. Data are presented as mean ± SD (n = 3).

### High Throughput Screening and Hit Identification

Based on the TAT-GESV validation experiments, we established a NanoBiT-based high-throughput screening workflow as illustrated in Figure 3, incorporating 3-hour compound incubation, appropriate controls, and counter-screening for luciferase interference. Using this optimized workflow, we proceeded to screen a diverse chemical library consisting of 10,240 compounds arranged in a 4-in-1 pooled DDSD plate format, where four structurally unrelated compounds were combined in each well to increase screening efficiency while maintaining adequate compound concentration for hit detection.

**Figure 3.**
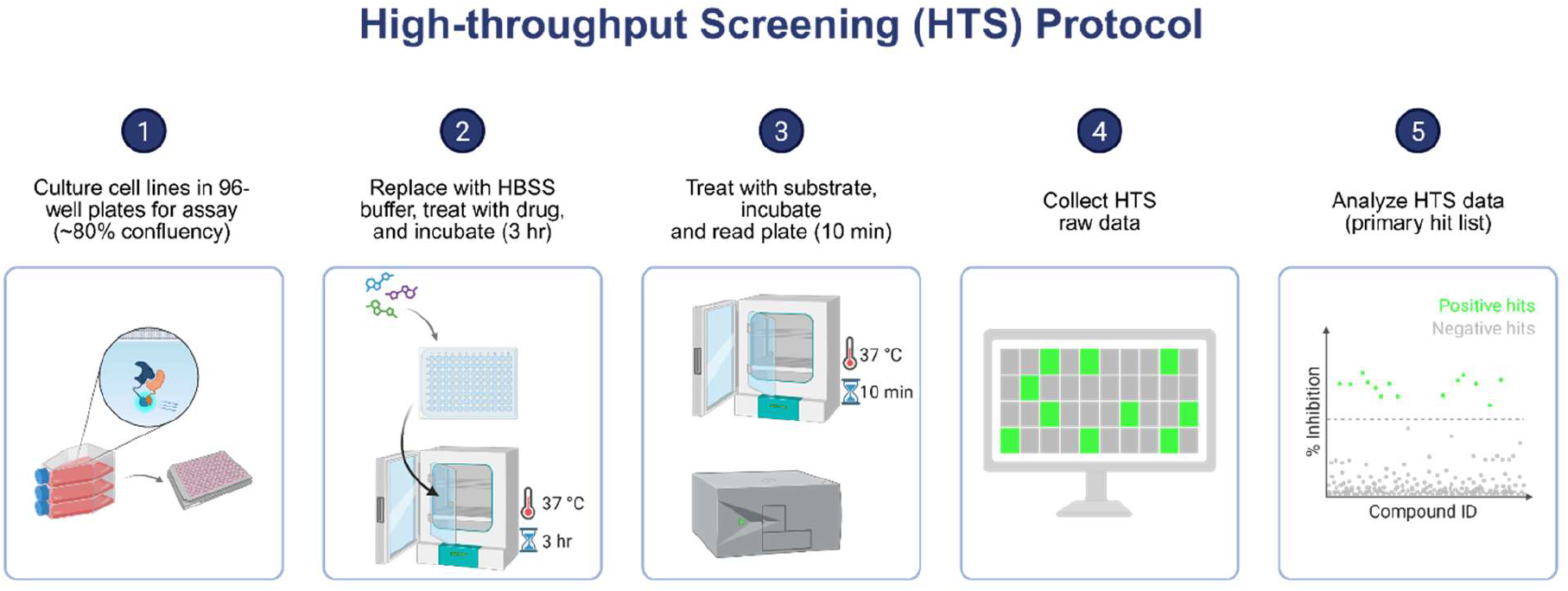
Our established workflow for the NOS1/NOS1AP NanoBiT assay

The screening workflow is summarized in Figure 4A. As shown in the heat map representation (Figure 4B), the primary screen at 25 μM compound concentration (total concentration of pooled compounds) yielded 54 hit groups showing ≥60% inhibition of the NanoBiT signal. To eliminate false positives arising from direct interference with NanoLuciferase activity, we performed a counter-screen using cells expressing only the reconstituted NanoLuciferase enzyme. This counter-screen identified 29 compounds that directly affected luciferase activity independent of the NOS1-NOS1AP interaction, and these were excluded from further analysis. The remaining 25 hit groups in the 4-in-1 format required deconvolution to identify the active single compounds. Each individual compound from these pooled groups was tested separately at 30 μM in the NanoBiT assay (Figure 4C). Single-compound validation confirmed 19 compounds as reproducible hits, with these compounds showing inhibition below 40% of control at 30 μM (green dots). This corresponds to a confirmed hit rate of 0.19% (19 hits from 10,240 compounds), which is consistent with typical hit rates observed in HTS campaigns targeting protein-protein interactions [20-22].

**Figure 4.**
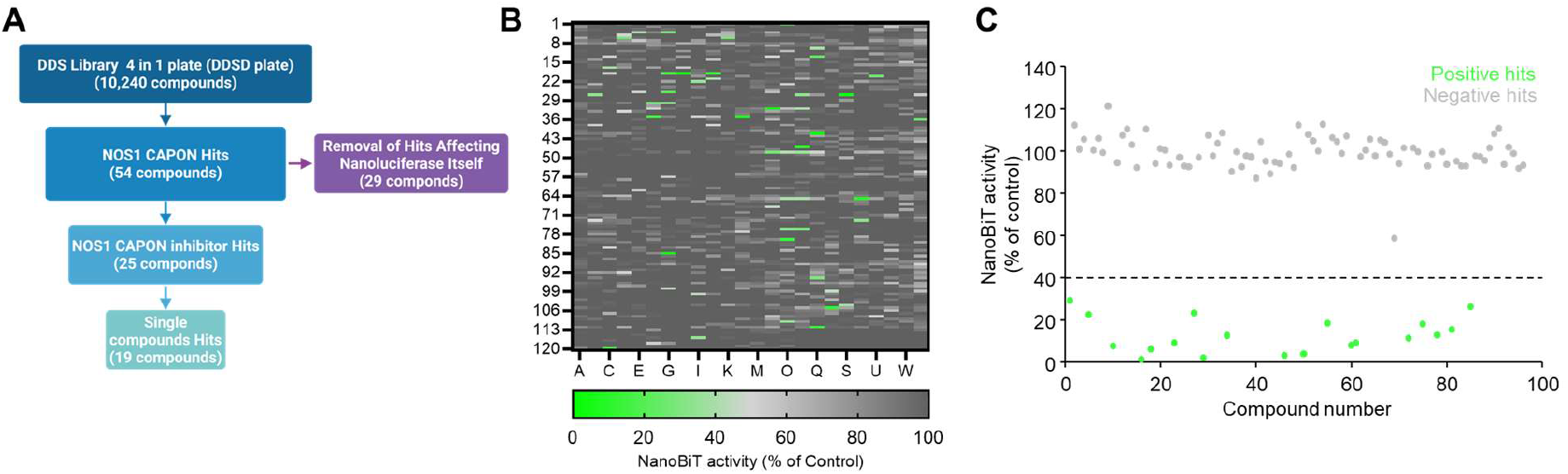
High-throughput screening for NOS1-NOS1AP interaction inhibitors using NanoBiT assay. Schematic diagram of the screening workflow. A total of 10,240 compounds from the DDS library (4 in 1 plate format) were screened, yielding 54 initial hits. After excluding 29 compounds that directly affected NanoLuciferase activity, 25 NOS1-NOS1AP inhibitor hits were identified. Subsequent single compound validation confirmed 19 hits. Heat map representation of primary screening results from 4 in 1 plates. NanoBiT activity is shown as percentage of control, with green indicating strong inhibition. (C) Validation of hits using single compounds at 30 μM. Green dots represent positive hits showing inhibition below 40% of control, while gray dots indicate negative hits.

### Characterization of hit compounds

To confirm that the 19 validated hit compounds inhibit the NOS1-NOS1AP interaction in a concentration-dependent manner rather than at a single fixed dose, we subjected these compounds to full dose-response analysis and successfully determined their IC_50_ values (Table 1). The compounds exhibited a range of potencies, with IC_50_ values ranging from 2.54 μM to greater than 30 μM. Notably, several compounds demonstrated sub-5 μM potency, including CN-4 (3.73 ± 0.37 μM), CN-8 (3.48 ± 0.07 μM), and CN-10 (2.54 ± 0.06 μM). These results indicate successful identification of multiple chemically distinct scaffolds capable of disrupting the NOS1-NOS1AP interaction with micromolar potency. We further characterized the three most potent compounds by examining their detailed dose-response profiles (Figure 5). The chemical diversity among these top hits suggests multiple binding modes may be accessible for inhibiting the NOS1-NOS1AP interaction, providing valuable starting points for medicinal chemistry optimization and structure-activity relationship studies.

**Table 1.** IC_50_ values of validated NOS1-NOS1AP interaction inhibitors. IC_50_ values were determined using the NanoBiT assay in CHO-K1 cells stably expressing NOS1-HiBiT and NOS1AP-LgBiT. Data are presented as mean ± SD (n = 3).

| Name | IC<sub>50</sub> (μM) | Name | IC<sub>50</sub> (μM) |
| --- | --- | --- | --- |
| CN-1 | 11.81 ± 5.36 | CN-11 | 4.97 ± 1.92 |
| CN-2 | 7.63 ± 3.73 | CN-12 | 24.92 ± 8.01 |
| CN-3 | 5.62 ± 0.13 | CN-13 | 9.68 ± 0.48 |
| CN-4 | 3.73 ± 0.37 | CN-14 | 5.82 ± 0.42 |
| CN-5 | 5.33 ± 0.16 | CN-15 | > 30 |
| CN-6 | 7.51 ± 0.16 | CN-16 | 6.63 ± 0.17 |
| CN-7 | 21.46 ± 4.89 | CN-17 | 6.98 ± 0.14 |
| CN-8 | 3.48 ± 0.07 | CN-18 | 9.33 ± 2.44 |
| CN-9 | 6.41 ± 0.45 | CN-19 | 6.8 ± 0.92 |
| CN-10 | 2.54 ± 0.06 | | |

**Figure 5.**
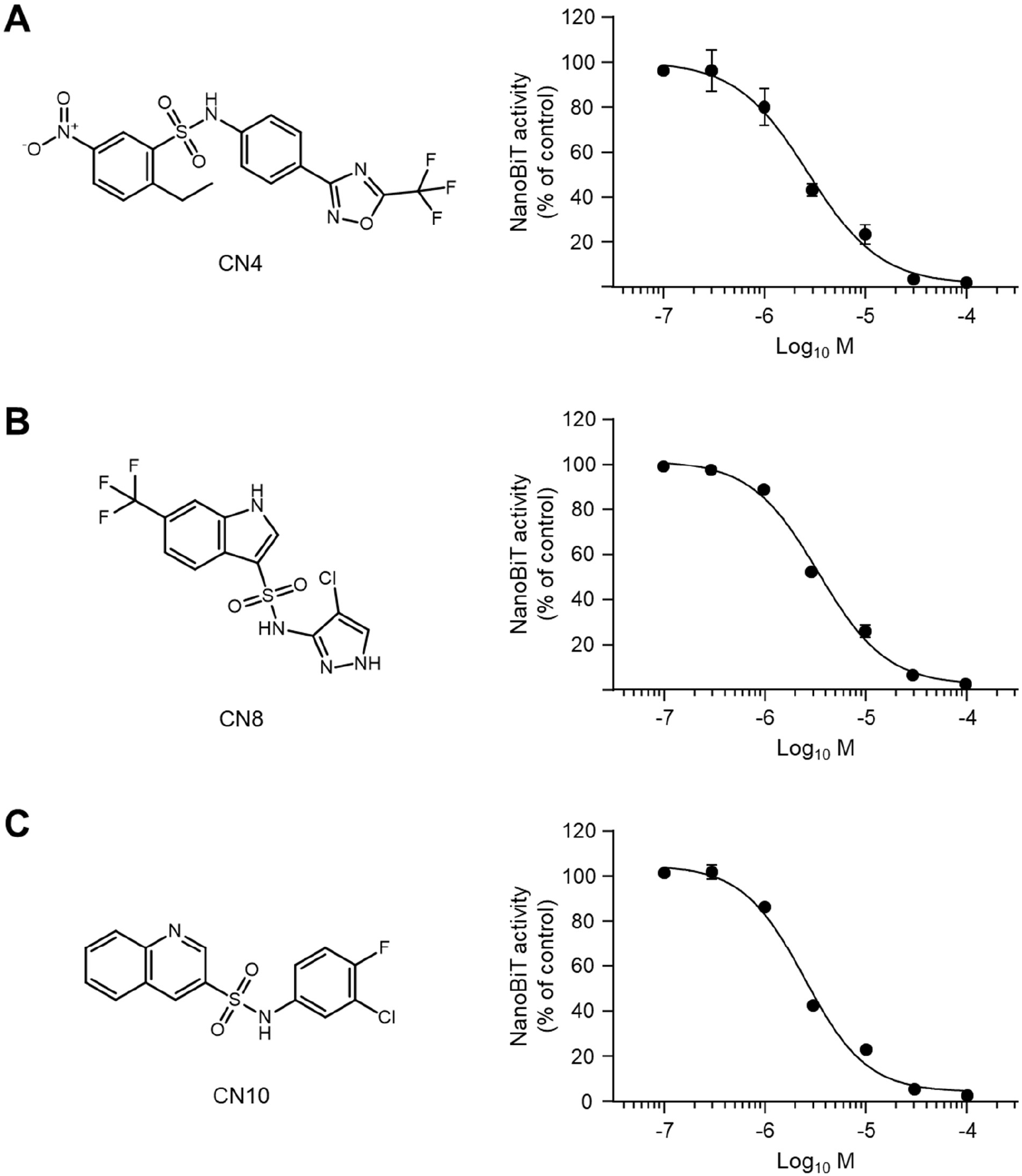
Chemical structures and dose-response curves of top three hit compounds. Chemical structures and concentration-dependent inhibition profiles of the three most potent compounds identified from the screen: (A) CN4, (B) CN8, and (C) CN10. NanoBiT activity is expressed as a percentage of vehicle control. Dose-response curves were generated from at least three independent experiments. Data are presented as mean ± SD (n = 3).

## Discussion

In this study, we successfully developed and validated a NanoBiT-based luminescence complementation assay for high-throughput screening of NOS1-NOS1AP interaction inhibitors. The assay demonstrated excellent performance characteristics, including a high signal-to-background ratio exceeding 240-fold and the ability to identify novel small molecule inhibitors with micromolar potency. This platform addresses a significant gap in the field, as previous methods for studying this protein-protein interaction were not well-suited for large-scale compound screening.

The NanoBiT technology offered several advantages over alternative approaches for our application. Compared to traditional FRET or BRET systems, NanoBiT provides superior brightness and stability, enabling more sensitive detection of protein interactions with lower background signal [14]. We specifically selected HiBiT rather than the original SmBiT for our NOS1 fusion protein based on several considerations. While SmBiT (K_d_ = 190 μM) was designed to minimize autonomous complementation and ensure that signal generation is driven entirely by the protein-protein interaction of interest, HiBiT (K_d_ = 700 pM) offers significantly higher affinity that translates to more robust signal generation in stable cell line systems [14, 15]. These technical advantages enabled us to establish a robust screening platform with excellent assay performance.

Using this optimized system, our NanoBiT based HTS campaign successfully identified 19 validated hits from a library of 10,240 compounds, yielding a hit rate of approximately 0.19%. This hit rate is consistent with typical protein-protein interaction screening campaigns, which generally yield hit rates of 0.01-1% depending on the assay system and compound library composition [20-23]. It has been reported that HTS campaigns targeting PPIs (protein-protein interaction) often yield lower hit rates compared to traditional enzyme targets due to the challenging nature of PPI interfaces [24]. The validation criteria employed in our study, including counter-screening to eliminate compounds that directly interfere with luciferase activity, ensures that the identified hits represent NOS1-NOS1AP interaction inhibitors rather than assay artifacts.

The IC_50_ values obtained for our hit compounds represent promising starting points for further optimization. While these potencies are modest compared to highly optimized drug candidates, they are typical for initial hits from phenotypic screens targeting protein-protein interactions [23, 25]. The NOS1-NOS1AP interface presents a particularly challenging target due to its relatively large contact surface area involving the PDZ domain and multiple interaction determinants [19, 26]. Nevertheless, several of our hits demonstrated sub-5 μM activity, suggesting that optimization through medicinal chemistry approaches could yield more potent analogs suitable for in vivo studies.

Such optimized compounds would be particularly valuable given the broad therapeutic relevance of the NOS1-NOS1AP interaction. In the cardiovascular system, NOS1AP genetic variants have been associated with QT interval prolongation and increased risk of sudden cardiac death, and modulation of its interaction with NOS1 could potentially offer therapeutic benefits in cardiac arrhythmias [5-7, 27]. In the central nervous system, disrupting this interaction has shown promise in preclinical models of excitotoxic neuronal death, and NOS1AP has been identified as a susceptibility gene for schizophrenia and other psychiatric disorders [8-11]. The compounds identified in our screen represent valuable tools for further investigating the therapeutic potential of targeting this protein-protein interaction in these disease contexts.

Despite these promising findings, our study has some limitations that should be acknowledged. First, the assay was performed in CHO-K1 cells, which may not fully recapitulate the native cellular environment of neurons or cardiomyocytes where NOS1-NOS1AP interactions are physiologically relevant. While CHO-K1 cells provided excellent assay performance and are widely used in drug discovery, future validation studies should examine compound activity in more disease-relevant cell types such as primary neurons or iPSC-derived cardiomyocytes. Second, the current assay format does not distinguish between compounds that directly prevent protein-protein interaction and those that might act through indirect mechanisms such as altering protein expression or stability. Furthermore, while we have confirmed disruption of the NOS1-NOS1AP interaction, further investigation is required to determine the binding site of these inhibitors— whether they interact with NOS1, NOS1AP, or potentially both proteins. Such target identification studies will be critical for rational optimization of these hit compounds.

To address these limitations and advance the characterization of identified hit compounds, we plan to conduct comprehensive biophysical studies in future work. Specifically, surface plasmon resonance (SPR) and microscale thermophoresis (MST) analyses using purified NOS1 and NOS1AP proteins will enable us to definitively determine which protein serves as the direct binding target for each inhibitor. These biophysical approaches will also provide quantitative binding affinity data (Kd values) and kinetic parameters (kon, koff) that are essential for understanding the mechanism of inhibition. Furthermore, differential scanning fluorimetry (DSF) or thermal shift assays could be employed to assess whether compounds induce conformational changes or affect protein stability. For compounds that demonstrate direct binding to either NOS1 or NOS1AP, structural studies using X-ray crystallography or cryo-electron microscopy may be pursued to elucidate the precise binding mode and guide structure-based optimization efforts. These future studies will be essential for advancing our most promising hit compounds toward therapeutic development.

In conclusion, we have established a robust NanoBiT-based HTS platform for identifying NOS1-NOS1AP interaction inhibitors and successfully applied it to discover 19 validated hit compounds, including three with sub-5 μM potency. This assay provides a valuable tool for the drug discovery community and represents a significant advancement in the ability to screen large compound libraries for modulators of this therapeutically relevant protein-protein interaction. The hits identified in this study serve as important starting points for medicinal chemistry optimization and further biological investigation, potentially leading to novel therapeutic agents for cardiovascular and neuropsychiatric disorders.

## CRediT authorship contribution statement

**Sungwoo Cho**: Writing – original draft, Methodology, Investigation, Data curation.

**Moustafa T. Gabr**: Writing – review & editing, Writing – original draft, Supervision, Project administration, Funding acquisition.

## Acknowledgements

This work was supported by the National Institute on Aging under grant number RF1AG084635-(PI: Gabr). We thank Dr. Katarzyna Kuncewicz for her assistance with the synthesis of the TAT-GESV peptide.

